# A novel pharmacogenetic testing panel for *CYP2C19* genetic polymorphisms

**DOI:** 10.64898/2026.02.14.705357

**Authors:** Miracle Nmesomachi Enwere, Rachelle Turiello, James Harrison McElroy, Elizabeth Medearis, Delaney Smith, Niklas Laurell, Renna Nouwairi, Anita H. Clayton, Atmaram Yarlagadda, Jerome C. Foo, Katherine J. Aitchison, B. Jill Venton, James P. Landers

**Author notes:** Co-senior authors (Jerome C. Foo, Katherine J. Aitchison, B. Jill Venton, James P. Landers).

## Abstract

Specific drug metabolism rates are defined by the constituency of the cytochrome P450 (CYP) genome, including polymorphic changes in any of 200+ CYP genes. An example is *CYP2C19,* where associations of gene polymorphisms with variability in certain drug metabolism rates have been linked to inter-individual and inter-ethnic differences in therapeutic outcomes. While pharmacogenomic screening for these variants prior to drug and dosage prescription has well-defined links to better treatment outcomes, current implementation is limited to complex and costly variant-probing and DNA sequencing protocols, which have limited availability in clinical laboratories, leading to slow turnaround times, impacting effective clinical intervention. Here we describe a novel, cost-effective, multiplex genotyping approach to screening *CYP2C19* variants. Fluorescence nested allele-specific (FAS) PCR was used with primers to detect *CYP2C19* variants of interest in specific ‘hot spots’, including the Tier 1 haplotypes identified by the Association for Molecular Pathology (AMP): *CYP2C19*2*, *\*3*, and *\*17.* The presence/absence of wild-type and mutant alleles were identified independently as haplotypes, and in a multiplex reaction as diplotypes representing the 10 possible genotype combinations/profiles. FAS-PCR achieved the same genotype calls as a pyrosequencing protocol optimized for validating genotypes, but with a simpler and more sensitive interface. The FAS-PCR method correctly identified the genotypes of both synthesized DNA and a human genomic DNA standard. Uniquely, the FAS-PCR protocol generates patterns using one fluorescently-labeled primer irrespective of the number of variants targeted, establishing it as considerably more cost-effective than other allele-specific PCR-based techniques that involve labeling both the forward and reverse primers.

## 1.0 INTRODUCTION

CYP2C19 is a P450 enzyme involved in the metabolism of a multiplicity of drugs, including many antidepressants, some cardiac drugs (such as clopidogrel, carisoprodol, and mavacamten), and some anticonvulsants (diazepam, brivaracetam, clobazam), and many others.^1–3^ It is encoded by the *CYP2C19* gene, which is located on chromosome 10q23.33 and spans 93.9kb.^4^ According to the Association for Molecular Pathology (AMP),^5^ three haplotypes of the *CYP2C19* gene are classified as Tier1 haplotypes because of “their well-established functional effect on CYP2C19 activity and drug response, availability of reference materials (RMs), and their frequencies in major ethnic groups.”^5^ These *CYP2C19* Tier 1 haplotypes are *CYP2C19*2*, *CYP2C19*3*, and *CYP2C19*17*.^5,6^ The main variant characterizing the *CYP2C19*2* (rs4244285) haplotype is a G to A substitution in location 681 (c.681G>A) of exon 5, which produces a cryptic splice site sequence that is preferentially selected, leading to the biosynthesis of a truncated, non-functioning CYP2C19 enzyme.^7^ The first functional variant in the *CYP2C19*3* haplotype to be identified was rs4986893, a c.636G>A in exon 4 that introduces a premature stop codon, leading to no functional CYP2C19 enzyme.^8^ From a functional perspective, the *CYP2C19*2* and *\*3* nucleotide substitutions are loss-of-function variants. The main gain-of-function variant in the *CYP2C19*17* haplotype is in the promoter region (c.-806C>T), and it increases the expression of CYP2C19 by increasing transcription factor and RNA polymerase binding during transcription.^4,9,10,11^ The PharmGKB Gene-Specific Information Tables, which support the Clinical Pharmacogenetics Implementation Consortium (CPIC) guidelines^12–14,15^ have a functional classification of *CYP2C19* Tier 1 diplotypes as follows: normal metabolizers (*\*1/*1*), poor metabolizers (*\*2/*2, *2/*3, *3/*3*), intermediate metabolizers (*\*1/*2* and *\*1/*3*), likely intermediate metabolizers (*\*2/*17* and *\*3/*17*), rapid metabolizers (*\*1/*17*), and ultrarapid metabolizers (*\*17/*17*).^9,10^ As the *CYP2C19* gene shares sequence similarity with other genes in the region, a sensitive assay that detects the mutations under consideration with a low false negative rate is essential. ^16^

The existing SNP genotyping techniques employed for screening the *CYP2C19* gene include, but are not limited to, bead-based immunoassay testing (e.g., xTAG^®^ CYP2C19 Kit v3),^16–18^ allele-specific polymerase chain reaction (PCR),^19,20^ fluorescence-based PCR (e.g., Spartan RX),^21^ next generation sequencing (NGS),^16,22^ Polymerase Chain Reaction-Restriction Fragment Length Polymorphism (PCR-RFLP),^7,23^ high-resolution melting (HRM) analysis,^24,25^ gene chip microarray technology (e.g., Affymetrix),^17,26,27^ and TaqMan probe chemistry.^16,28^ Most, if not all, of these technologies rely on *in vitro* hybridization of single-stranded oligonucleotides (primers or probes) to a target DNA sequence associated with the gene(s) of interest. Genotype discrimination power relies on perfectly matched or mismatched target-probe/primer hybridization with oligonucleotides immobilized on a solid support, e.g., bead-based immunoassay testing, gene chip microarray technology^29^ and NGS ( e.g., Illumina sequencing);^30,31^ or free in a buffered solution, as with HRM analysis, PCR-RFLP, allele-specific PCR, among others. The use of multi-parallel arrays that carry out solid support detection is notable in that high throughput and multiplexed genotyping can be achieved. Pyrosequencing has also been employed to assess *CYP2C19*2*, *\*3*, and *\*17* allele frequencies in a Korean population.^32^ However, these techniques requires costly reagents and equipment that most testing laboratories have not widely adopted. Additionally, bioinformatics expertise is crucial for processing and analyzing computationally intensive high-throughput data generated by NGS techniques.^33^ PCR-RFLP has the limitation of only being able to detect mutations that occur within restriction endonuclease splice sites. It’s time-consuming nature is also considered a major disadvantage. A relatively economic method is TaqMan via an OpenArray, in multiple assays for genes, including *CYP2C19* may be multiplexed together for multiple samples. However, if one is interested only in *CYP2C19*, much of the assays would be redundant.

Here, we describe a fluorescent nested allele-specific PCR (FAS-PCR) methodology, which is based on the variable fragment length allele-specific polymerase chain reaction (VFLASP) technique by Tóth et al.^34^ for the generation of a multiplex pharmacogenetic panel for genotyping the *CYP2C19* Tier 1 haplotypes. Synthesized DNA representing wild-type and mutant-type versions of the *CYP2C19*2, *3,* and *\*17* haplotypes were tested and validated using pyrosequencing. The system was also challenged with a commercial standard of deidentified human genomic DNA, with a demonstration of excellent sensitivity for detecting SNPs, with data being easy to interpret from both qualitative (presence or absence of peaks/allele) and quantitative (DNA size comparison) standpoints.

## 2.0 METHODS

### 2.1 Study design and sample materials

This study was approved by the University of Virginia (UVA) Institutional Review Board (IRB) under the UVA Center for Psychiatric Clinical Research. Two classes of DNA samples were used for this study: Synthesized DNA representing wild-type and mutant-type mimics of the *CYP2C19*2, *3,* and *\*17* alleles (Integrated DNA Technologies or IDT, Coralville, Iowa, USA) and deidentified human genomic DNA commercialized by Promega (Promega Corporation, Madison, WI 53711 USA). The primary sequence of the IDT-synthesized DNA can be found in **Figures S1-S6**.

The IDT-synthesized DNA (known as gBlocks) was ordered to include flanking regions of the *CYP2C19*2* (NC_000010.11:g.94781721-94781858<rs4244285<NC_000010.11:g.94781860-94782140), *\*3* (NC_000010.11:g.94780461-94780652<rs4986893<NC_000010.11:g.94780652-94780880), and *\*17* (NC_000010.11:g.94761746-94761899<rs12248560<NC_000010.11:g.94761901-94762123) haplotypes. The synthesized DNA was 420 bp in size for the *\*2* and *\*3* haplotypes and 378 bp for the *\*17* haplotypes. The disparity in the size of synthesized DNA was based on the allowable complexity of the DNA fragment defined by the IDT. Both the WT and MT of the haplotypes under consideration were synthesized, making a total of six positive controls. The Promega DNA was used as the test sample. The IDT-synthesized DNA of known genotype was used for assay validation.

Using the six positive controls, we set up three detection systems: singleplex, multiplex, and pyrosequencing systems, as seen in **Figure 1** and explained in the sections that follow.

**Figure 1.**
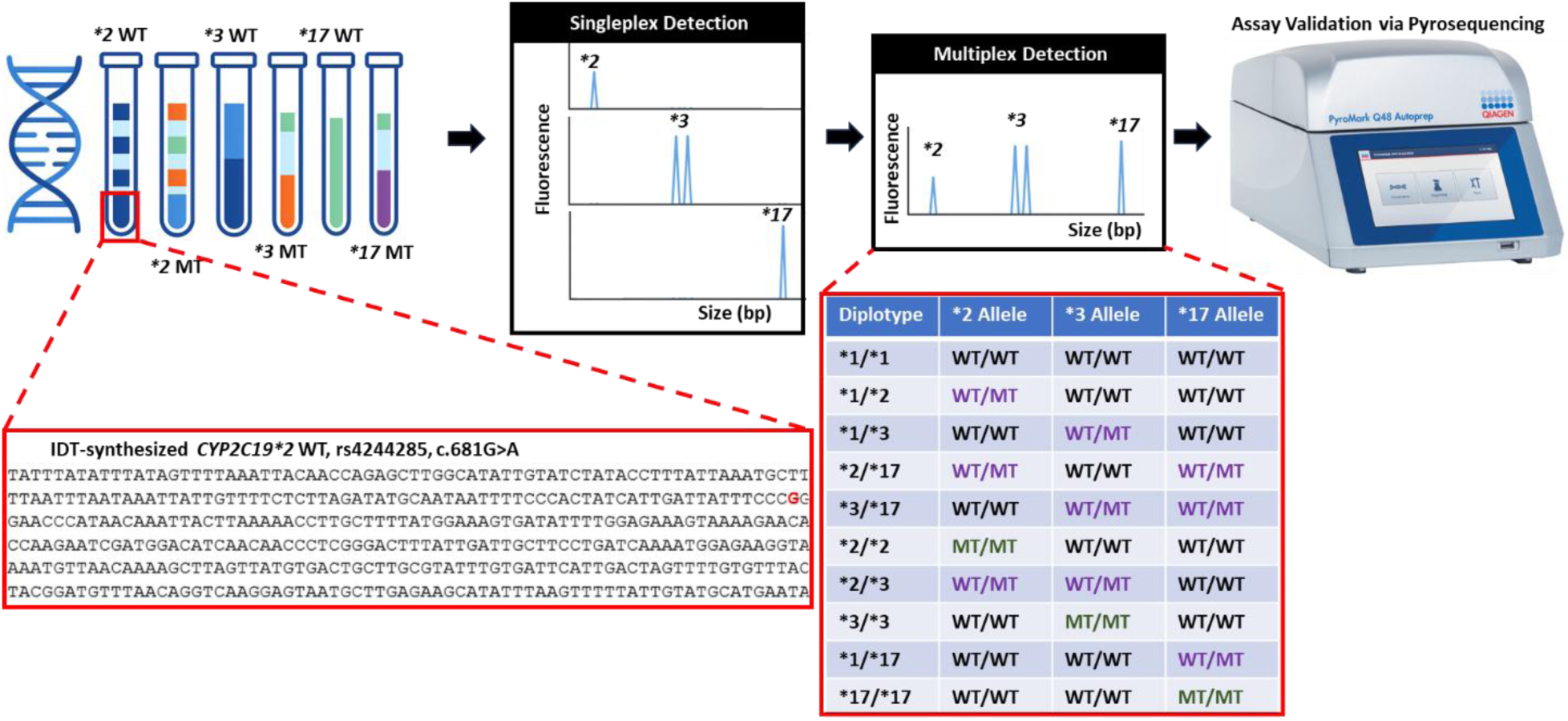
Workflow for the development of the FAS-PCR technique, where the arrows show. Six synthesized DNA positive controls representing wild-type and mutant-type of *CYP2C19*2, *3,* and *\*17* haplotypes have known primary sequence. A singleplex detection platform was set up to assess the allele-specific functionality of the primers for detecting wild-type and/or mutant types of the haplotypes under consideration at each locus separately. A multiplex detection platform was set up to assess the allele-specific functionality of the primers for simultaneous single-tube detection of all possible diplotypes as presented in the table. The result of the singleplex setup is validated by pyrosequencing, which cannot be used for multiplex allele detection. The pyrosequencing data indirectly validate the already known primary sequence of all positive controls.

#### 2.2.1 Protocol for singleplex detection

The first step in the development and design of the multiplex panel for *CYP2C19*2, *3*, and *\*17* haplotypes was to assess each haplotype separately. This was done to assess whether or not the synthesized primers worked for their intended purpose.

The PCR reactions were completed using a Veriti™ 96-Well Fast Thermal Cycler in a total volume of a 25 μL reaction, with 2 μL DNA template (25ng human genomic DNA) or 1 μL of 10 ng positive controls (IDT-synthesized DNA), 2mM MgCl_2_, 200 μM deoxyribonucleotide phosphate (dNTP) mixture, 2.5 μL of 10x PCR Buffer II (Life Technologies), 7.5μL of nuclease-free water, 5μL of 1.0 U AmpliTaq Gold DNA Polymerase (Life Technologies, Carlsbad, CA), 0.2 μM of M13(− 21) 5′-FAM labeled primer, 0.2 μM of both allele-specific forward primers with adapters, and 0.04 μM of M13(−21) 5′ adapter sequence linked reverse primer. PCR conditions were optimized to comprise an initial denaturation step of 95℃ for 10 min, followed by 30 cycles of denaturation at 95°C for 20 s, annealing at 55°C for 30 s, and extension at 72°C for 45 s, followed by a final extension cycle at 72°C for 10 min. All primers were synthesized by the Integrated DNA Technologies, IDT (Coralville, Iowa, USA).

In a single PCR system, we added SNP-specific forward primers in equimolar concentrations to the FAM-labeled M13(−21) primer. The concentration of the M13(−21) 5’ reverse primer was lower because it is only involved in the initial amplification step from which PCR amplicons are yielded. Afterward, the FAM-labeled M13(−21) primer, which was added in excess, binds only to amplicons that contain the reverse complement of the M13(−21) sequence (originally from the reverse primer).

#### 2.2.2 Protocol for multiplex detection

Having established the viability of the primers, the second step in the development and design of the multiplex panel for *CYP2C19*2, *3*, and *\*17* haplotypes was to assess all haplotypes in a single tube.

The PCR reactions were completed using a Veriti™ 96-Well Fast Thermal Cycler in a calculated volume of 25 μL, including 3 μL DNA template (1 μL or 0.5 μL of 10 ng positive controls as specified in **Table 1** or 1.5 μL of 25ng of test sample), 2 mM MgCl_2_, 200 μM dNTP mixture, 2.5 μL of 10x PCR Buffer II (Life Technologies), 11.4 μL of nuclease-free water, 0.8 μL of 2.0 U AmpliTaq Gold DNA Polymerase (Life Technologies, Carlsbad, CA). For ease of reference, the optimized primer concentrations are given in **Table S4**. PCR conditions were optimized to comprise an initial denaturation step of 95°C for 10 min, followed by 30 cycles of denaturation at 95°C for 20 s, annealing at 55°C for 30s, and extension at 72°C for 45 s, followed by a final extension cycle at 72°C for 10 min.

**Table 1.** Classification Based on Metabolizer Status.

| **Diplotype** | **\*2 Locus** | **\*3 Locus** | **\*17 Locus** | **Metabolizer Status** |
| --- | --- | --- | --- | --- |
| *\*1/\*1* | WT/WT | WT/WT | WT/WT | Normal |
| *\*1/\*2* | WT/MT | WT/WT | WT/WT | Intermediate |
| *\*1/\*3* | WT/WT | WT/MT | WT/WT | Intermediate |
| *\*2/\*17* | WT/MT | WT/WT | WT/MT | Likely Intermediate |
| *\*3/\*17* | WT/WT | WT/MT | WT/MT | Likely Intermediate |
| *\*2/\*2* | MT/MT | WT/WT | WT/WT | Poor |
| *\*2/\*3* | WT/MT | WT/MT | WT/WT | Poor |
| *\*3/\*3* | WT/WT | MT/MT | WT/WT | Poor |
| *\*1/\*17* | WT/WT | WT/WT | WT/MT | Rapid |
| *\*17/\*17* | WT/WT | WT/WT | MT/MT | Ultrarapid |

WT: wild-type haplotype; MT: mutant-type haplotype; WT/WT: wild-type haplotypes in both maternal and paternal chromosomes; MT/MT: mutant-type haplotypes in both maternal and paternal chromosomes; WT+ MT: wild-type and mutant-type haplotypes in maternal and paternal chromosomes, respectively.
WT/WT contained 1 µL of the WT DNA associated with a locus. It simulates the wild-type nucleotide in both maternal and paternal chromosomes (Homozygous wild-type diplotype).
MT/MT contained 1 µL of the MT DNA associated with a locus. It simulates the mutant-type nucleotide in both maternal and paternal chromosomes (Homozygous mutant-type diplotype).
WT/MT contained 0.5 µL WT and 0.5 µL MT of the DNA associated with a locus. It simulates the wild-type and mutant-type nucleotide in either maternal or paternal chromosome (Heterozygous diplotype).

Using the algorithm (genotype/diplotype to phenotype translation) in **Table 1**, the positive controls (450bp wild-type and mutant-type gene fragments covering the *2, *3, and *17 loci) were grouped into their specific metabolizer status. For the algorithm, we employed the Pharmacogenomics Knowledgebase (PharmGKB) diplotype to phenotype translation system for *CYP2C19* by extracting all the possible diplotype combinations of the three haplotypes (*\*2, *3,* and *\*17*).^35^ This correlates with the diplotypes indirectly described by Kim et al.^32^ in an attempt to classify *CYP2C19* polymorphisms in a multiplex pyrosequencing panel. The hypothesis was that the human genomic DNA test sample would fall under one of the diplotypes or their corresponding metabolizer statuses described.

### 2.3 Electrophoresis

For the separation, detection, and analysis of DNA fragments, two capillary electrophoresis (CE) instruments were employed: the 2100 Bioanalyzer (Agilent Technologies, Santa Clara, CA, USA) and the Spectrum Compact CE System (CE1304 Promega Corporation, Madison, WI 53711 USA). The 2100 Bioanalyzer was used for all PCR optimization protocols since it does not require cost-intensive fluorescently labeled amplicons for the detection of fragments. However, due to its ±5 bp peak resolution,^36^ we employed the Spectrum Compact CE instrument that has single base pair resolution for the multiplex FAS-PCR protocol. The two instruments were used according to manufacturer recommendations. For the 2100 Bioanalyzer, 9 µL of prepared gel dye matrix in a light-protected (amber) tube, 5 µL of DNA 1000 marker (green cap), and 1 µL of DNA 1000 ladder were used from the Agilent DNA 1000 kit (Agilent Technologies, Santa Clara, CA, USA). In addition to 1 µL of PCR amplicons, the reagents in the Agilent DNA 1000 kit were added to their respective wells in the DNA microfluidic chip (Agilent Technologies, Santa Clara, CA, USA). Fragment analysis was then completed using the 2100 Expert software (Agilent Technologies, Santa Clara, CA, USA).

The FAM-labeled amplicons were separated and detected using the Spectrum Compact CE instrument and analyzed using GeneMarker HID v.2.8.2 (SoftGenetics, LLC, State College, PA, USA). Each well contained 9.5 µL of Hi-Di™ formamide (Applied Biosystems™, Waltham, MA, USA), 0.5 µL of WEN Internal Lane Standard (ILS) 500 (Promega), and 1 µL of DNA amplicon.

### 2.4 Pyrosequencing validation study for screening the *CYP2C19*2*, *\*3*, and *\*17* loci

To confirm if the results from the FAS-PCR technique were accurate, we employed a genotyping technique known as pyrosequencing. The pyrosequencing primers were the same as those of Kim et al.^32^ However, instrumentation and protocol differed. This is because the pyrosequencing step was conducted according to manufacturer instructions using the PyroMark Q48 Autoprep Instrument and Q48 Advanced Reagents. The PyroMark Q48 Autoprep Instrument Software Version 4.3.3 (QIAGEN, Germantown, MD 20874, USA) was used to visualize, generate, and analyze all pyrosequencing data in this study.

Since the PyroMark Q48 Autoprep Instrument (QIAGEN, Germantown, MD 20874, USA) used is not capable of multiplex sequencing, the loci under consideration (*\*2*, *\*3*, *\*17*) had to be screened separately. The pyrosequencing step involved the use of two reagent kits: the PyroMark PCR kits (QIAGEN, Germantown, MD 20874, USA) and the Q48 Advanced Reagents (Qiagen), for the PCR and pyrosequencing steps, respectively.

The pyrosequencing PCR reactions were completed using a Veriti™ 96-Well Fast Thermal Cycler in a total volume of a 25 μL, with 2 μL DNA template [1μL WT (10ng) and 1μL MT (10ng) of *\*2, *3,* and *\*17* IDT-synthesized DNA for heterozygous genotypes, 2μL of either WT (10ng) or MT (10ng) of *\*2, *3,* and *\*17* IDT-synthesized DNA for homozygous genotypes, or 2 μL of 25ng of test sample], 7μL of nuclease-free water, 10X CoralLoad Concentrate (QIAGEN, Germantown, MD 20874, USA), 0.2 μM of (biotinylated) forward primer (for *\*2*, *\*3*, or *\*17* screening), 0.2 μM of (biotinylated) reverse primer (for *\*2*, *\*3*, or *\*17* screening), and 0.04 μM of Pyromark PCR Master Mix (QIAGEN, Germantown, MD 20874, USA). PCR conditions were the same as the multiplex FAS-PCR protocol outlined above. The forward primers for screening the *\*2* and *\*3* loci were biotinylated at the 5′ end whereas only the reverse primer of the *\*17* locus was biotinylated according to the design of Kim et al.^32^

## 3.0 RESULTS AND DISCUSSION

### 3.1 Assay Design and Development

The allele-specific PCR (AS-PCR) core component of the assay facilitates the specific binding, elongation, and amplification of primers that have a nucleotide at the 3’-end that is complementary to the SNP of interest. The allele-specific primers designed by Rehman *et al*. ^37^ were leveraged to screen the *CYP2C19*2* locus, while the primers for screening the *CYP2C19*3* and *\*17* loci were custom-designed using NCBI BLAST.^38^ Rehman *et al.*^37^ employed AS-PCR to assess the allelic distribution of the *CYP2C19*2* variants in Pakistani cardiac patients receiving clopidogrel treatment. However, to improve the selectivity of the primers, three changes were made here: 1) a special adapter known as a Pigtail (GTTTCTTT) was added to the 5′-end of the forward primers, 2) an M13(−21) sequence was added to the 5′-end of the reverse primers, and 3) a 3 bp variable sequence was also incorporated in the forward primer that targets the mutant (A) SNP as specified in **Table S1**.

The Pigtailed sequence was added to the 5’-end of the forward SNP-specific (wild-type and mutant-type) primers because this modification has been shown in the literature^39,40^ and experimental analysis involving the screening of the *CYP2C19*2* locus (**Figure S7**), to increase primer selectivity and reduce the level of stutter observed with downstream electrophoresis. The primers without Pigtailed sequence showed stutter bands while the primers with the Pigtailed sequence showed high selectivity, as seen by the lack of stutter. In this way, the conventional AS-PCR protocol, which was tested with and without the adapter sequences, was optimized to increase selectivity for the *CYP2C19*2* region of interest upon incorporation of these adapters (**Figure S1**). Therefore, all the forward primers (for *\*2*, *\*3*, and *\*17* screening) were designed with the Pigtailed sequence.

The M13(−21) sequence incorporation is essential for the generation of the reverse complement of the M13(−21) sequence for the effective binding of fluorescein (FAM)-labeled M13(−21) primer in a nested PCR phase (**Figures 2 and 3**). This step is crucial since it yields fluorescently labeled amplicons.

**Figure 2.**
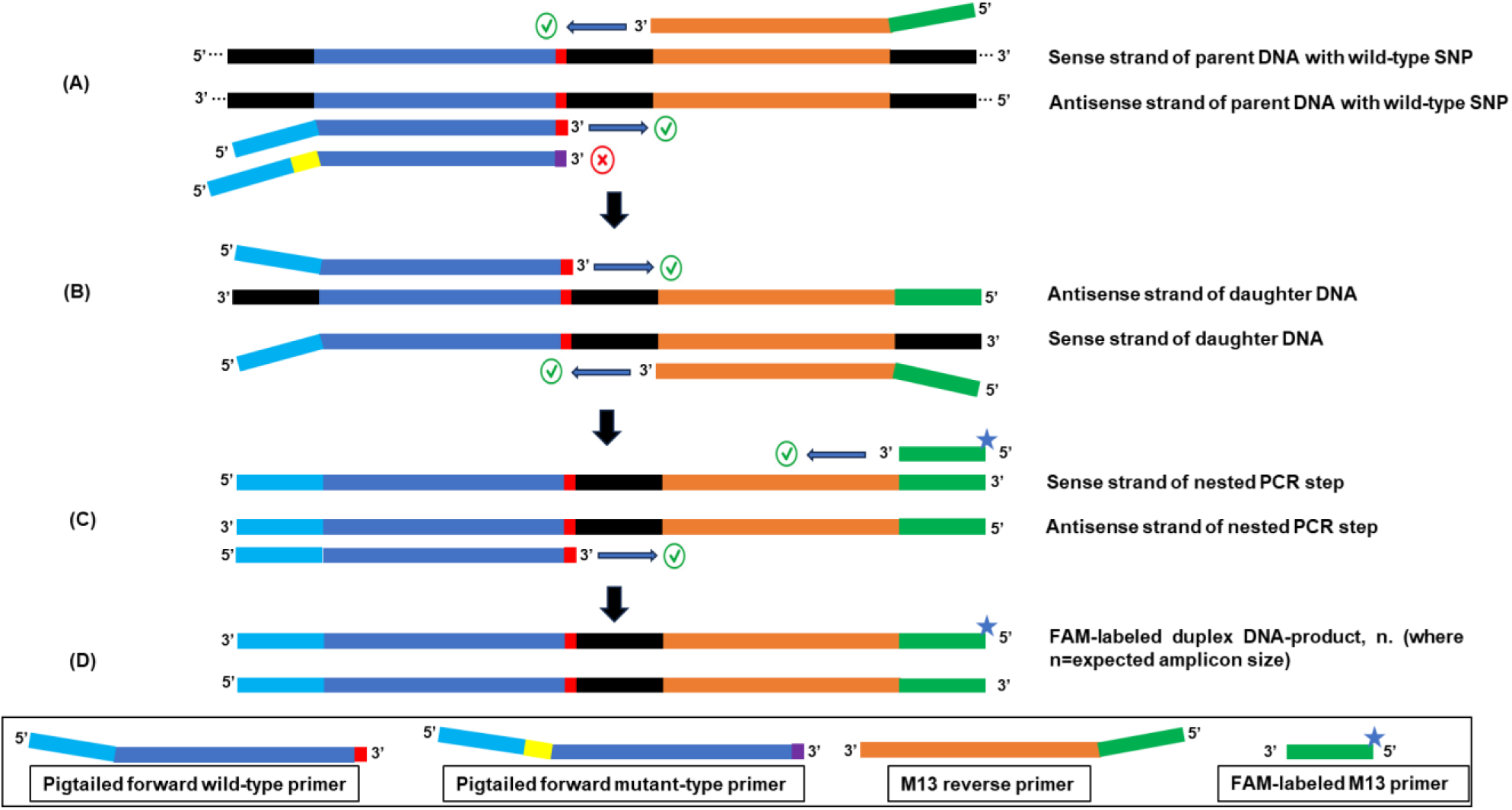
Allele-specific amplification of wild-type DNA sequence. The red color represents the wild-type SNP of interest whereas purple represents the mutant-type SNP **(A)** Allele-specific primers anneal to their complementary regions on the parent DNA (DNA template). Only primers complementary to the SNP of interest at their 3’ end are successfully amplified by DNA polymerase. **(B)** A nested PCR step ensues, which is characterized by amplification of PCR products from the initial step by complementary allele-specific primers. **(C)** Fluorescein (FAM)-labeled M13 primer binds only to its complementary sequence from the nested PCR step. PCR amplification then takes place. **(D)** The duplex DNA product formed is fluorescently labeled.

**Figure 3.**
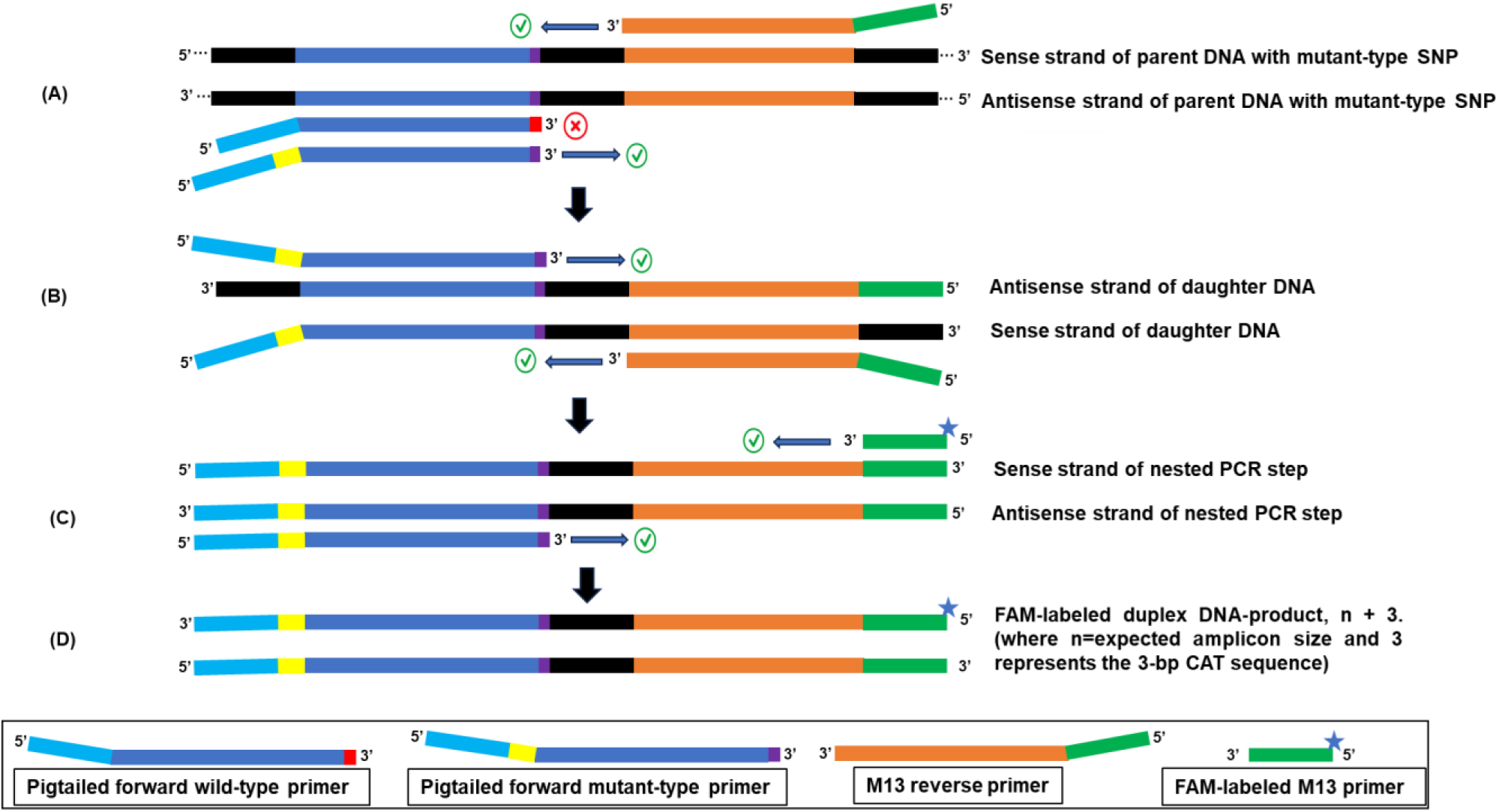
Allele-specific amplification of mutant-type DNA sequence. The purple color represents the mutant SNP of interest whereas red represents the wild-type SNP **(A)** Allele-specific primers anneal to their complementary regions on the parent DNA (DNA template). Only primers complementary to the SNP of interest at their 3’ end are successfully amplified by DNA polymerase. (**B)** A nested PCR step ensues which is characterized by amplification of PCR products from the initial step by complementary allele-specific primers. **(C)** Fluorescein (FAM)-labeled M13 primer binds only to its complementary sequence from the nested PCR phase. PCR amplification then takes place. **(D)** The duplex DNA product formed is fluorescently labeled.

The third modification was made to further distinguish amplicons containing the wild-type (WT) nucleotide (**Figure 2**) from those containing the mutant-type (MT) nucleotide (**Figure 3**) after separation and detection by electrophoresis. Specifically, the forward SNP primer that selectively amplifies the mutant haplotype was designed so that a 3-base pair (bp) CAT sequence (not complementary to the template DNA) was sandwiched between the Pigtailed sequence and complement of the target sequence on the forward primer. This innovation distinguishes heterozygous populations of a haplotype for which each haplotype is visualized as two distinct bands separated by 3 bp in the resultant electropherogram. This design was used to reduce cost and complexity, as there was no need to fluorescently label the wild-type and mutant-type (*CYP2C19*2, *3,* and *\*17*) haplotypes in this study. Instead of labeling all six forward primers that target the wild-type and mutant-type haplotypes with fluorophores that are expensive, the FAS-PCR technique involves fluorescently labeling one primer: the M13(−21) primer.

### 3.2 Validation of Positive Controls by Pyrosequencing

Pyrosequencing was used to validate our samples and FAS-PCR protocol. We wanted to first confirm that the synthesized DNA positive controls met the quality standards for the development of our FAS-PCR assay. Pyrosequencing was therefore used to genotype the *CYP2C19*2*, *\*3*, and *\*17* loci independently. The genotype profile for each locus is represented in **Figure 4**. In the top panel of Figures **4A**, **4B**, and **4C**, we show normal metabolizer (*\*1/*1*) genotypes of the *\*2, *3,* and *\*17* haplotypes, respectively. These are seen as single (or dominant) peaks corresponding to the presence of the WT haplotype. In the middle panel of each of **Figures 4A**, **4B**, and **4C**, intermediate (*\*1/*2*, and *\*1/*3*) and rapid metabolizer (*\*1/*17*) status are shown by two peaks of similar percentage heights of the wild-type peak heights corresponding to the WT and MT nucleotides of the *\*2, *3*, and *\*17* haplotypes. Poor (*\*2/*2* and *\*3/*3*) and ultrarapid (*\*17/*17*) metabolizer status is seen as single or dominant peaks corresponding to the presence of the MT haplotypes in the lower panel of **Figure 4**. Peak heights less than 20% were deemed negligible when compared with the negative controls, and these may be due to background noise.

**Figure 4.**
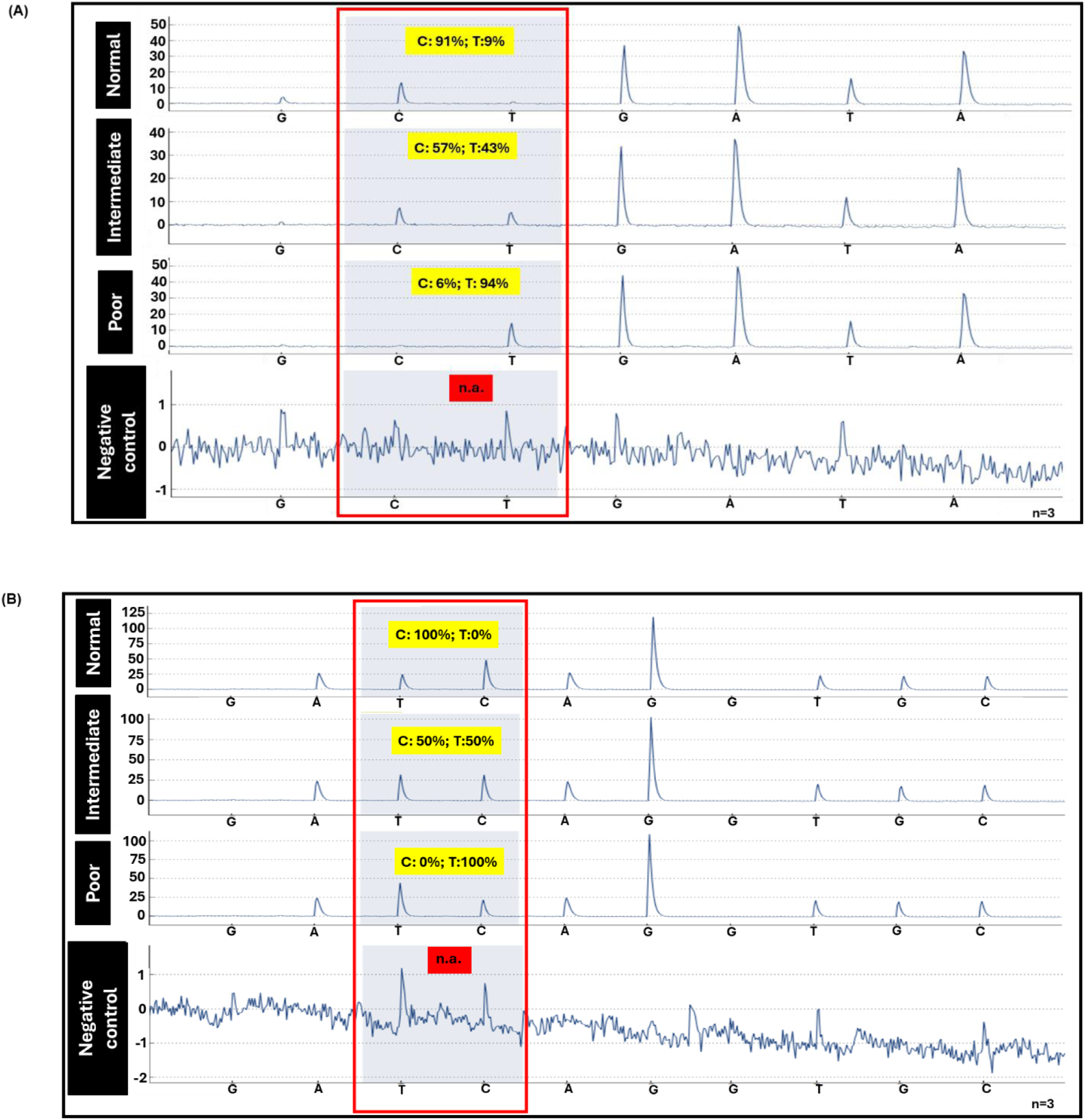
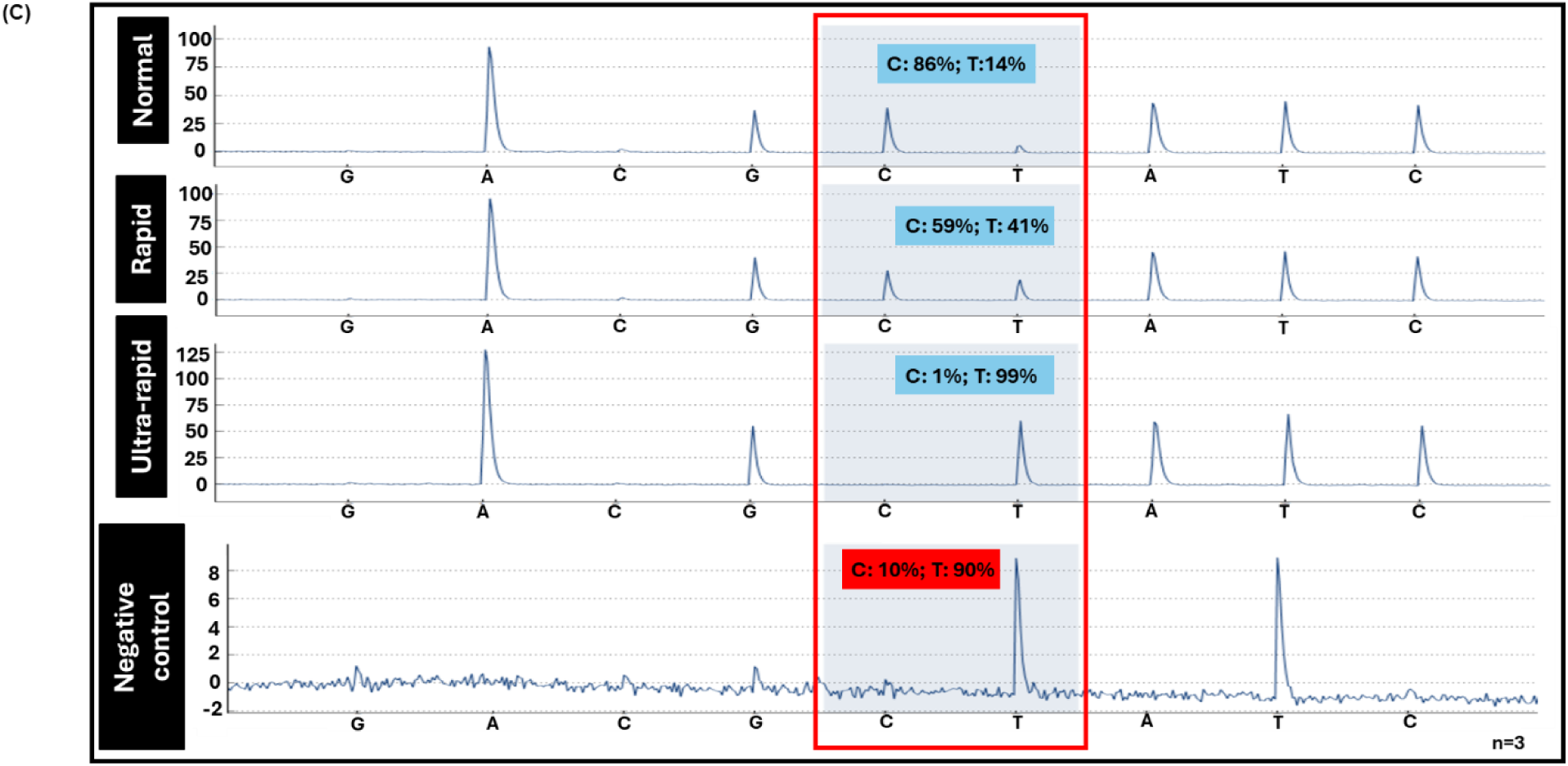
Representative pyrosequencing electropherograms (pyrograms) for CYP2C19-SNP profiles and their associated metabolizer status as predicted by pyrosequencing software. Regions of interest are marked by the large red boxes. The x-axis is the sequence analyzed and the y-axis represents bioluminescence intensity. **(A)** The SNP profile of the *CYP2C19*2* locus and its associated metabolizer status. Homozygous (single) peaks for normal and poor metabolizers are characteristically represented by nucleotides C and T, respectively. Intermediate metabolizer status is heterozygous and can be detected by the panel with two distinct peaks corresponding to C and T. **(B)** The SNP profile of the *CYP2C19*3* locus and its associated metabolism status. Homozygous (single) peaks for normal and poor metabolizers are characterized by the highest peaks which correspond to C and T (the complement of G and A), respectively. Intermediate metabolizer status is heterozygous and can be detected by the panel with two distinct peaks of equal heights. **(C)** The SNP profile of the *CYP2C19*17* locus and its associated SSRI-metabolism status. Homozygous (single) peaks for normal and ultra-rapid metabolizers are characteristically represented by nucleotides C and T, respectively. Intermediate metabolizer status is heterozygous can be detected by the panel with two distinct peaks corresponding to C and T. All samples and negative controls were run in replicates of three (n=3).

### 3.3 FAS-PCR Genotyping

#### 3.3.1 Singleplex Detection

After confirming the quality of the IDT-synthesized DNA samples, we went on to test our FAS-PCR protocol. We began testing the FAS-PCR primers for screening each haplotype (*\*2*, *\*3*, and *\*17*) independently for singleplex detection. The data can be interpreted both qualitatively (based on relative locus-specific peak height ratio cut-off) and quantitatively (based on amplicon size). As shown in **Figure 5**, the successful representation of heterozygous genotypes (*\*1/*2, *1/*3,* and *\*1/*17*) was characterized by two peaks that are 3 bp away from each other. Homozygous WT DNA samples were indicated by a single peak at approximately 152 bp, 189 bp, and 253 bp, corresponding to WT nucleotide at the *CYP2C19*2*, *\*3*, and *\*17* loci, respectively. These loci were screened independently, i.e., in separate reaction systems, and all have the *\*1/*1* designation. In the same manner, homozygous MT DNA samples were associated with a single electrophoretic peak at a position that is 3 bp greater than that for the homozygous WT. The sizes of the homozygous MT genotypes are approximately 155 bp, 192 bp, and 256 bp which correspond to the detected MT nucleotide at the *CYP2C19*2* (*\*2/*2*), *\*3* (*\*3/*3*), and the *\*17* (*\*17/*17*) loci, respectively. These results are the same as from pyrosequencing and confirm that FAS-PCR primers targeting all three (*\*2, *3*, and *17) loci were ready to be tested in a multiplex reaction.

**Figure 5.**
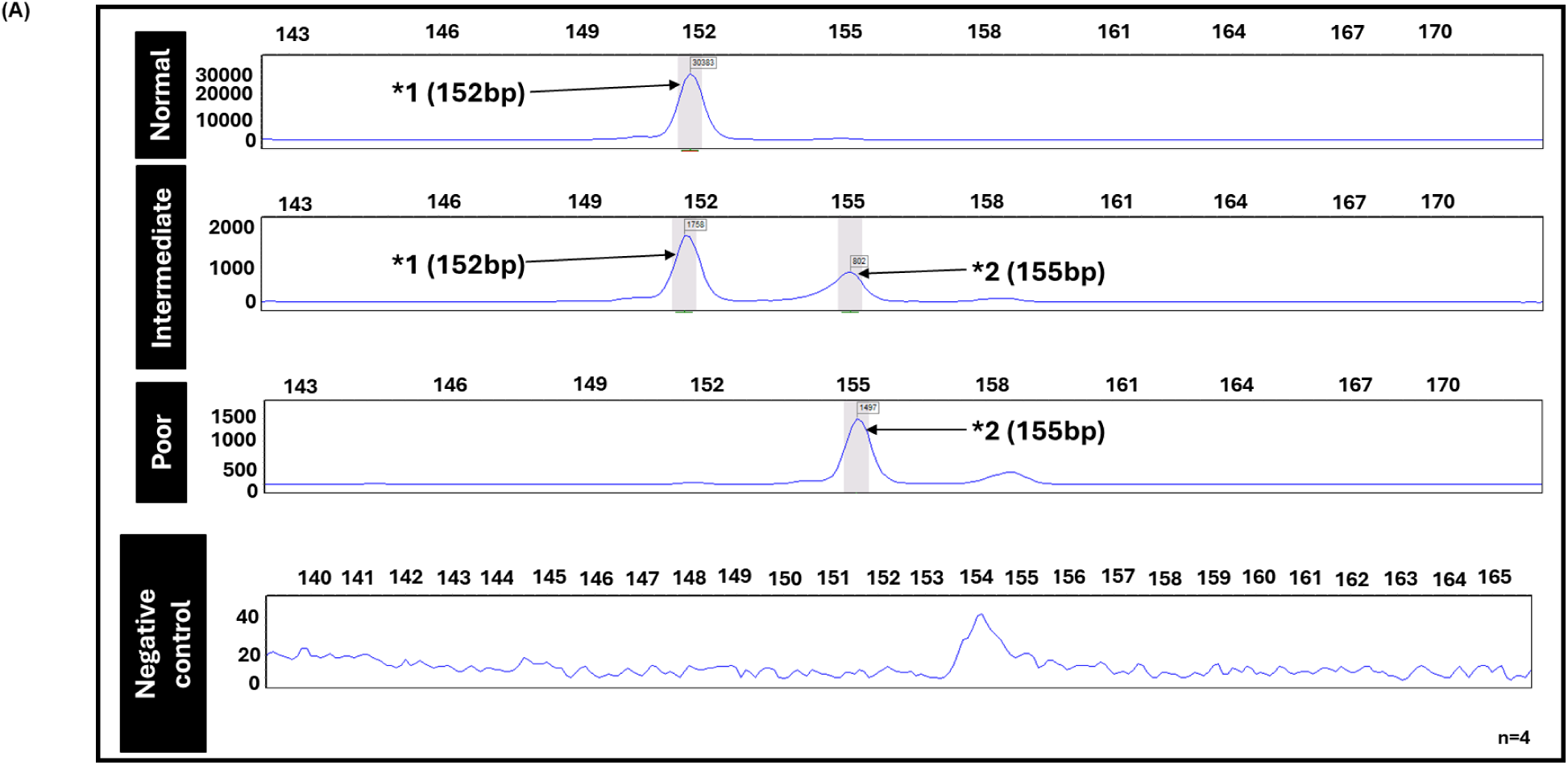
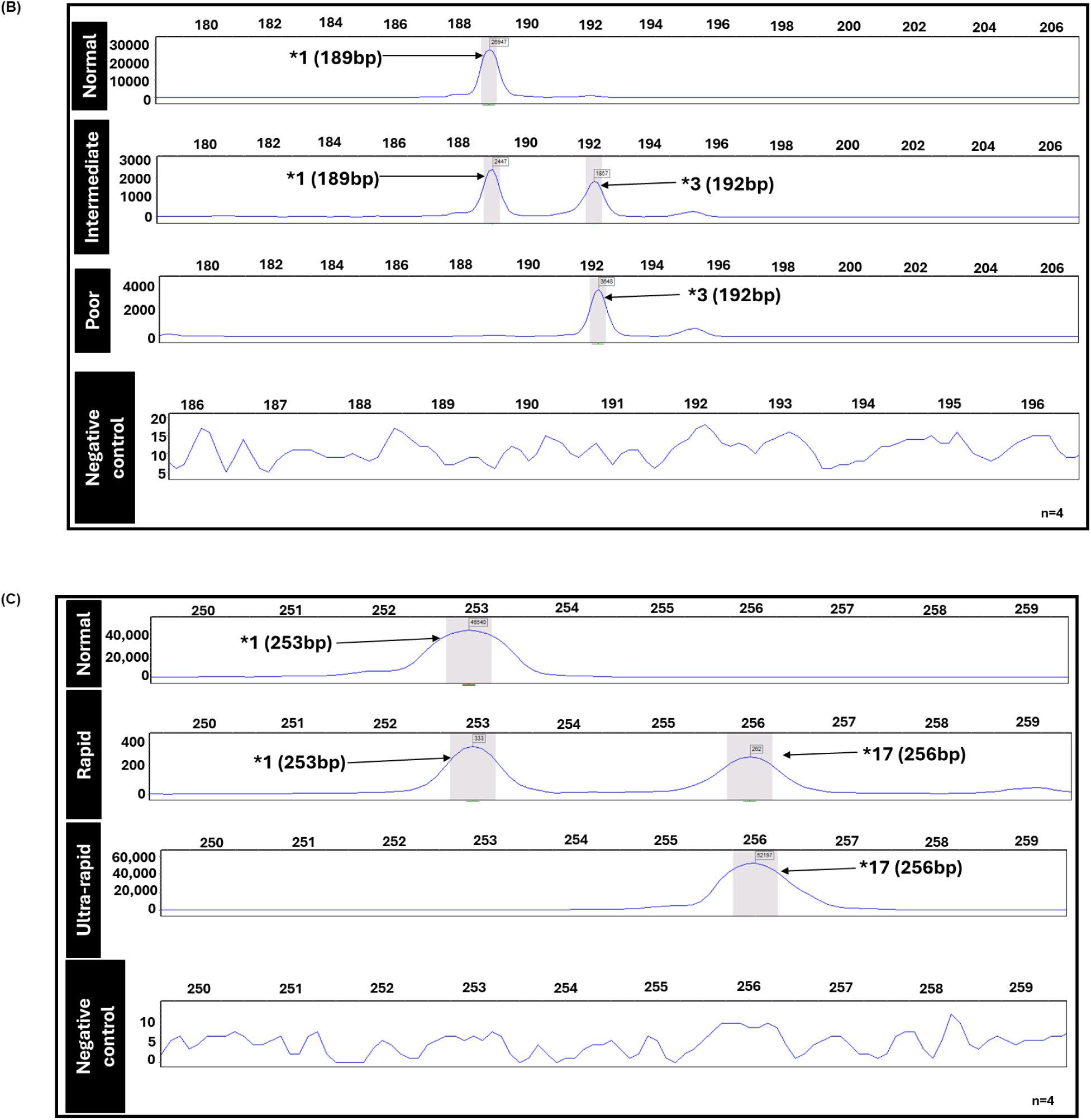
Representative electropherograms showing *CYP2C19*-SNP profiles and their associated metabolizer status. The x-axis is the fluorescence intensity and the y-axis represents DNA fragment size in base-pairs (bp). (A) The SNP profile of the *CYP2C19*2* locus and its associated SSRI-metabolism status. Homozygous (single) peaks for normal and poor metabolizers are characteristically 152 bp and 155 bp in length, respectively. Intermediate metabolizer status is heterozygous can be detected by the panel with two distinct peaks that are 152 bp and 155 bp in length. **(B)** The SNP profile of the *CYP2C19*3* locus and its associated SSRI-metabolism status. Homozygous (single) peaks for normal and poor metabolizers are characteristically 189 bp and 192 bp in length, respectively. Intermediate metabolizer status is heterozygous can be detected by the panel with two distinct peaks that are 189 bp and 192 bp in length. **(C)** The SNP profile of the *CYP2C19*17* locus and its associated SSRI-metabolism status. Homozygous (single) peaks for normal and ultra-rapid metabolizers are characteristically 253 bp and 256 bp in length, respectively. Intermediate metabolizer status, is heterozygous can be detected by the panel with two distinct peaks that are 253 bp and 256 bp in length. All samples and negative controls were run in replicates of four (n=4).

#### 3.3.2 Multiplexed Detection

After ascertaining that the primers facilitate amplification in a standalone setup, a multiplex protocol was optimized with the primers needed to screen for all the genotypes (**Table 1**) in a single run. Thus, each tube had a master mix that contained forward and reverse primers for screening the WT and MT forms of the *CYP2C19*2*, *\*3*, and *\*17* haplotypes. The master mix also contained the reverse primers for all haplotypes. **Table 1** was simulated in-tube using IDT-synthesized DNA (**Figure 6**). The data can also be interpreted qualitatively and quantitatively as indicated in section 3.3.1. The genotypes were classified based on the relative peaks at the locations specified in section 3.3.1 for different combinations of WT and/or MT at specific *CYP2C19* loci.

**Figure 6.**
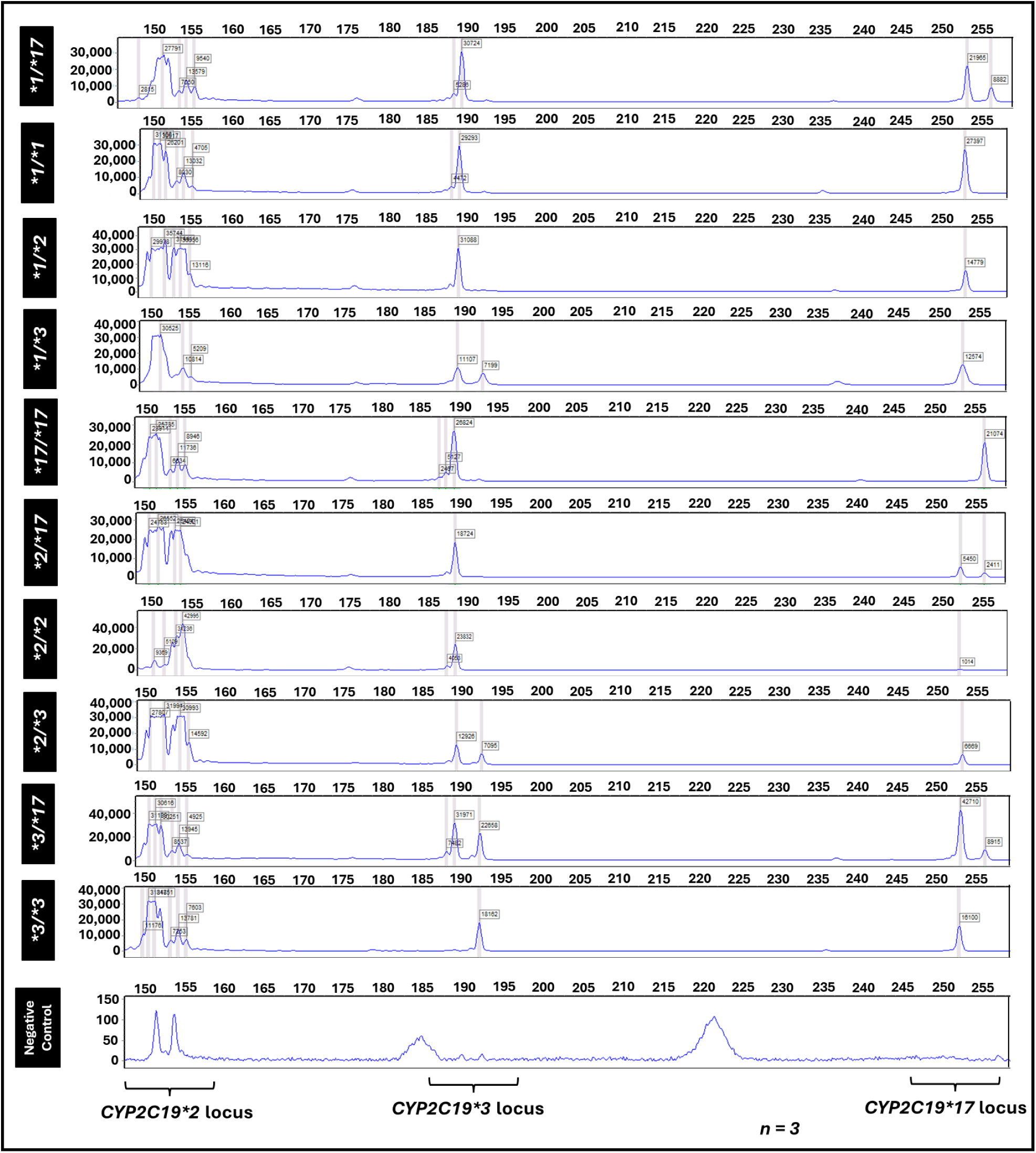
Representative electropherograms showing various *CYP2C19*-SNP profiles according to Tier1 status. The x-axis is the fluorescence intensity and the y-axis represents DNA fragment size in base-pairs (bp). The data can be interpreted both qualitatively (based on amplicon size). and quantitatively (based on relative locus-specific peak height). The *CYP2C19*2* locus shows either homozygous (single) peaks for wild-type (WT) and mutant-type (MT) haplotypes at 152 bp and 155 bp, respectively or heterozygous (two) distinct peaks that are 152 bp and 155 bp in length. The negative control spanning the *CYP2C19*2* locus suggests some contamination. The *CYP2C19*3* locus show either homozygous (single) peaks for wild-type (WT) and mutant-type (MT) haplotypes at 189 bp and 192 bp, respectively, or heterozygous (two) distinct peaks that are 189 bp and 192 bp in length. The *CYP2C19*17* locus shows either homozygous (single) peaks for wild-type (WT) and mutant-type (MT) haplotypes at 253 bp and 256 bp, respectively or heterozygous (two) distinct peaks that are 253 bp and 256 bp in length. All samples and negative controls were run in replicates of three (n=3).

### 3.4 Assessing FAS-PCR Performance using a Test Sample

The final step was to evaluate the performance of our FAS-PCR technique using extracted human genomic DNA. This process involved the generation of a FAS-PCR genetic profile from a test sample and subsequent validation of the result using pyrosequencing (**Figure 7**). The obtained FAS-PCR panel results indicated that the test sample had a *2/*17 genotype and is, this, an intermediate metabolizer; this result was validated by pyrosequencing. The red, green, and purple boxes show the screening result of the *CYP2C19*2*, *CYP2C19*3*, and *CYP2C19*17* loci, respectively. They indicate WT and MT for the *CYP2C19*2* locus, WT for the *CYP2C19*3* locus, and WT and MT for the *CYP2C19*17* locus.

**Figure 7.**
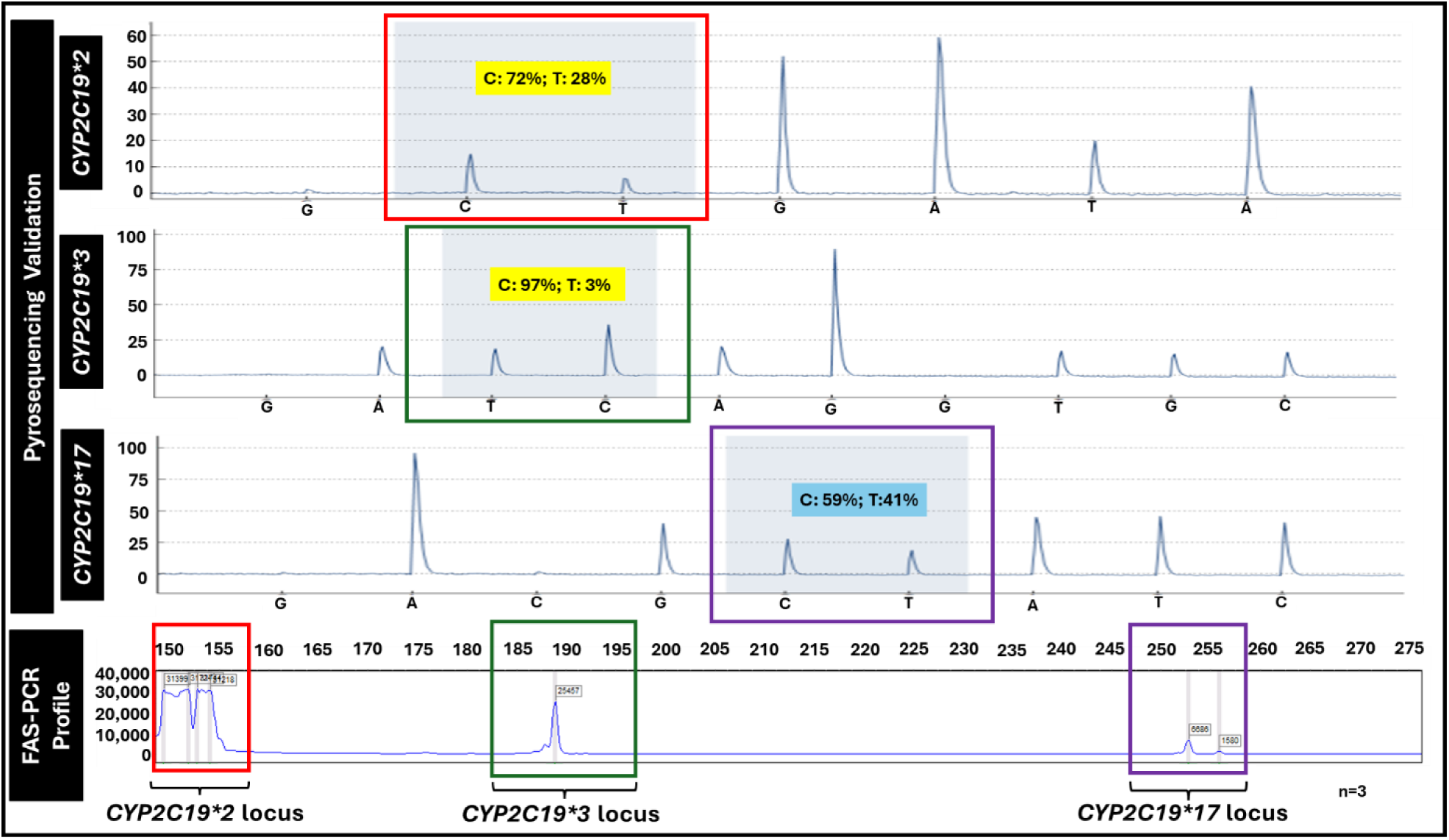
Representative electropherograms showing the screening of a test DNA sample using pyrosequencing and the FAS-PCR technique. The red, green, and purple boxes highlight the *CYP2C19*2*, *CYP2C19*3*, and *CYP2C19*17* locus, respectively. The pyrosequencing validation revealed that in the test sample screening of the *CYP2C19*2* region, both wild-type (WT) and mutant-type (MT) haplotypes at 72% and 28% were present. Also, the pyrogram **of the *CYP2C19*3* region** shows wild type peaks at 97% and a negligible mutant-type peak at 3%. The ***CYP2C19*17* locus** shows that both wild-type (WT) and mutant-type (MT) *17 haplotypes at 59% and 41% were present. The FAS-PCR profile panel shows heterozygosity (that is, WT and MT) for *CYP2C19*2* and *\*17* but homozygosity for *CYP2C19*3* WT. All samples were run in replicates of three (n=3).

## 4.0 CONCLUSION

Drugs are prescribed with the assumption that the consumer is a normal metabolizer, thus underserving the two extremes of metabolizers, ultrarapid and poor metabolizers, leading to adverse drug reactions, lack of efficacy, and potentially, death. For example, CYP2C19 poor metabolizers should receive half the standard starting dose of escitalopram or citalopram, otherwise they are at increased risk of corrected-QT (QTc) prolongation and potential sudden cardiac death.^42,43^ One of the reasons for the slow implementation of pre-emptive pharmacogenetic testing is that the commercially available kits are prohibitively expensive and time-consuming. Here, we reported a novel method of screening the *CYP2C19* gene for the *\*2*, *\*3*, and *\*17* haplotypes that predispose individuals to metabolize CYP2C19-mediated drugs differently. We developed fluorescent nested allele-specific PCR (FAS-PCR), an adaptation of the VFLASP, for rapid pharmacogenomic screening.^34^ It involves the single-tube amplification of SNPs in multiple genes to assess the presence or absence of a haplotype. The double stranded [is there a word missing here?] of fluorescently-labeled amplicons were separated and detected as single-stranded amplicons by capillary electrophoresis and laser-induced fluorescence detection, respectively. Our adaptation of the VFLASP involves multiplex detection of SNPs. A methodologically similar panel using the allele-specific technique by Cypa et al^44^ was found to be 10x cheaper, 5x faster, and comparable to targeted sequencing in terms of accuracy. Our novel FAS-PCR protocol has potential cost benefits over existing technologies and the pharmacogenomic panel of Cypa et al^44^ because it minimizes the use of expensive fluorophores through innovations in the primer design for allele discrimination. Genotypes were identified based on the size of either wild-type or mutant-type genotypes. Our novel method could potentially be used to analyze the DNA of patients to inform the dosage of other prescription drugs metabolized by the CYP2C19 enzyme, such as clopidogrel^45^ and warfarin.^46^

The limitations of this novel methodology include the following. During the singleplex and multiplex optimization protocol, determining a single PCR parameter that enables detection of WT or MT alleles of the haplotypes under consideration is a major challenge. There is usually a trade-off that allows non-specific amplifications in the electropherograms that are not within the expected amplicon sizes. In addition, further validation of this technique using human genomic DNA is required.

In addition to its utility for pharmacogenomic testing, the reported novel SNP genotyping method has a potentially broad range of applications. For example, Adebiyi and Enwere et al.^47^ have shown that the strains and variants of Severe acute respiratory syndrome coronavirus 2 (SARS-CoV-2) can be detected by identifying SNPs in specific genes using FAS-PCR?. Drug-resistant microbes and SNPs associated with genetics-based diseases, such as cancer, epilepsy, and multiple sclerosis, could also be detected using this technology. These applications present potential future directions for our novel methodology. Some foreseeable bottlenecks include optimization of the reported protocol due to the presence of PCR inhibitors such as ethanol. Further studies could also involve automation of the FAS-PCR technique for use in point-of-care settings. This new method of FAS-PCR gives insight into the rate of metabolism of drugs metabolized by the CYP2C19 enzyme and aids personalized treatment based on specific genotypes.

## Supporting information

Supplementary Table S1. Primers employed for genotyping of CYP2C19*2

## AUTHOR CONTRIBUTION STATEMENT

This manuscript was written by M.N.E., J.C.F., K.J.A., A.H.C., N.L., A.Y., B.J.V., and J.P.L. The research was designed by M.N.E., R.T., R.N., B.J.V., and J.P.L. The research was performed by M.N.E., J.H.E., E.M., and D.S. The data was analyzed by M.N.E., J.C.F., K.J.A., B.J.V., and J.P.L.

## Notes

### Competing Interest Statement

The authors have declared no competing interest.

### Funding

- National Institutes of Health (NIH): R01MH137467

