## Supplementary Table S1. Primers employed for genotyping of CYP2C19*2 for "A novel pharmacogenetic testing panel for *CYP2C19* genetic polymorphisms"

**Section 1. FAS-PCR Primer Information**

**Table S1.** Primers employed for genotyping of *CYP2C19*2*

| Primers | Primer Sequence |
| --- | --- |
| **PIGtailed_CYP_Fwd_WT (*2)** | 5'- **GTGTCTTCCACTATCATTGATTATTTCCCG** -3' |
| **PIGtailed_CYP_Fwd_MT (*2)** | 5'- **GTGTCTTCATCCACTATCATTGATTATTTCCCA** -3' |
| **M13(-21)_CYP_Rev (*2)** | 5'- **TGTAAAACGACGACGGCCAGTTAAAGTCCCGAGGGTTGTTG** -3' |
| **FAM-labeled_M13(-21)_primer** | 5'- /FAM/**TGTAAAACGACGACGGCCAGT**-3' |

**Table S2.** Primers employed for genotyping of *CYP2C19*3*

| Primers | Primer Sequence |
| --- | --- |
| **PIGtailed_CYP_Fwd_WT (*3)** | **5'-GTGTCTTGGATTGTAAGCACCCCCTGG-3'** |
| **PIGtailed_CYP_Fwd_MT (*3)** | 5'-**GTGTCTTCATGGATTGTAAGCACCCCCTGA**-3' |
| **M13(-21)_CYP_Rev (*3)** | 5'-**TGTAAAACGACGACGGCCAGTTTTCCAGATATTCACCCCATGG** -3' |

**Table S3.** Primers employed for genotyping of *CYP2C19*17*

| Primers | Primer Sequence |
| --- | --- |
| **PIGtailed_CYP_Fwd_WT(*17)** | **5'-GTGTCTTGTGTCTTCTGTTCTCAAAGC -3'** |
| **PIGtailed_CYP_Fwd_MT (*17)** | 5'-**GTGTCTTCATGTGTCTTCTGTTCTCAAAGT** -3' |
| **M13(-21)_CYP_Rev (*17)** | 5'-**TGTAAAACGACGACGGCCAGTTACCTTAGCAGATATAAACACCT**-3' |

*Nucleobases highlighted: Blue- PIGtail, Yellow-3bp discriminant, Green- M13(-21), Red- allele-specific nucleotide

**Table S4.** Primer Concentrations

| Reagent | Working Concentration | Final Concentration |
| --- | --- | --- |
| PIGtailed_CYP_Fwd_WT (*2) | $10\mu M$ | $0.2\mu M$ |
| PIGtailed_CYP_Fwd_WT(*3) | $10\mu M$ | $0.2\mu M$ |
| PIGtailed_CYP_Fwd_WT(*17) | $10\mu M$ | $0.2\mu M$ |
| PIGtailed_CYP_Fwd_MT (*2) | $10\mu M$ | $0.2\mu M$ |
| PIGtailed_CYP_Fwd_MT(*3) | $10\mu M$ | $0.2\mu M$ |
| PIGtailed_CYP_Fwd_MT(*17) | $10\mu M$ | $0.2\mu M$ |
| M13(-21)_CYP_Rev (*2) | $10\mu M$ | $0.08\mu M$ |
| M13(-21)_CYP_Rev (*3) | $10\mu M$ | $0.08\mu M$ |
| M13(-21)_CYP_Rev (*17) | $10\mu M$ | $0.08\mu M$ |
| FAM-labeled_M13(-21)_primer | $10\mu M$ | $0.6\mu M$ |

### Section 2. DNA Template Synthesized by the Integrated DNA Technologies (IDT)

```
>NC_000010.11:g.94781721-94782140 Homo sapiens chromosome 10,
GRCh38.p14 Primary Assembly
TATTTATATTTATAGTTTTAAATTACAACCAGAGCTTGGCATATTGTATCTATACCTTTATTAAATGCTT
TTAATTTAATAAATTATTGTTTTCTCTTAGATATGCAATAATTTTCCCACTATCATTGATTATTTCCCGG
GAACCCATAACAAATTACTTAAAAACCTTGCTTTTATGGAAAGTGATATTTTGGAGAAAGTAAAAGAACA
CCAAGAATCGATGGACATCAACAACCCTCGGGACTTTATTGATTGCTTCCTGATCAAAATGGAGAAGGTA
AAATGTTAACAAAAGCTTAGTTATGTGACTGCTTGCGTATTTGTGATTCATTGACTAGTTTTGTGTTTAC
TACGGATGTTTAACAGGTCAAGGAGTAATGCTTGAGAAGCATATTTAAGTTTTTATTGTATGCATGAATA
```

**Figure S1.** Primary Sequence of IDT-Synthesized DNA Wild-Type DNA template for CYP2C19*2

```
>NC_000010.11:g.94780461-94780880 Homo sapiens chromosome 10,
GRCh38.p14 Primary Assembly
TGTTTACTCATATTTTAAAATTGTTTCCAATCATTTAGCTTCACCCTGTGATCCCACTTTCATCCTGGGC
TGTGCTCCCTGCAATGTGATCTGCTCCATTATTTTCCAGAAACGTTTCGATTATAAAGATCAGCAATTTC
TTAACTTGATGGAAAAATTGAATGAAAACATCAGGATTGTAAGCACCCCCTGGATCCAGGTAAGGCCAAG
TTTTTTGCTTCCTGAGAAACCACTTACAGTCTTTTTTTCTGGGAAATCCAAAATTCTATATTGACCAAGC
CCTGAAGTACATTTTTGAATACTACAGTCTTGCCTAGACAGCCATGGGGTGAATATCTGGAAAAGATGGC
AAAGTTCTTTATTTTATGCACAGGAAATGAATATCCCAATATAGATCAGGCTTCTAAGCCCATTAGCTCC
```

**Figure S2.** Primary Sequence of IDT-Synthesized DNA Wild-Type DNA template for CYP2C19*3

```
>NC_000010.11:g.94761746-94762123 Homo sapiens chromosome 10,
GRCh38.p14 Primary Assembly
TAAGTGGTTCTATTTAATGTGAAGCCTGTTTTATGAACAGGATGAATGTGGTATATATTCAGAATAACTA
ATGTTTGGAAGTTGTTTTGTTTTGCTAAAACAAAGTTTTAGCAAACGATTTTTTTTTTCAAATTTGTGTC
TTCTGTTCTCAAAGCATCTCTGATGTAAGAGATAATGCGCCACGATGGGCATCAGAAGACCTCAGCTCAA
ATCCCAGTTCTGCCAGCTATGAGCTGTGTGGCACCAACAGGTGTCCTGTTCTCCCAGGGTCTCCCTTTTC
CCATTTGAAATATAAAAAATAACAATTCCTGCCTTCACGTGTTTTTTTAGGGGGTTAAATGGTAAAGGTG
TTTATATCTGCTAAGGTAATTTACTTGA
```

**Figure S3.** Primary Sequence of IDT-Synthesized DNA Wild-Type DNA template for CYP2C19*17

```
>NC_000010.11:g.94781721-94782140 Homo sapiens chromosome 10,
GRCh38.p14 Primary Assembly
TATTTATATTTATAGTTTTAAATTACAACCAGAGCTTGGCATATTGTATCTATACCTTTATTAAATGCTT
TTAATTTAATAAATTATTGTTTTCTCTTAGATATGCAATAATTTTCCCACTATCATTGATTATTTCCCAG
GAACCCATAACAAATTACTTAAAAACCTTGCTTTTATGGAAAGTGATATTTTGGAGAAAGTAAAAGAACA
CCAAGAATCGATGGACATCAACAACCCTCGGGACTTTATTGATTGCTTCCTGATCAAAATGGAGAAGGTA
AAATGTTAACAAAAGCTTAGTTATGTGACTGCTTGCGTATTTGTGATTCATTGACTAGTTTTGTGTTTAC
TACGGATGTTTAACAGGTCAAGGAGTAATGCTTGAGAAGCATATTTAAGTTTTTATTGTATGCATGAATA
```

**Figure S4.** Primary Sequence of IDT-Synthesized DNA Mutant-Type DNA template for CYP2C19*2

```
>NC_000010.11:g.94780461-94780880 Homo sapiens chromosome 10,
GRCh38.p14 Primary Assembly
TGTTTACTCATATTTTAAAATTGTTTCCAATCATTTAGCTTCACCCTGTGATCCCACTTTCATCCTGGGC
TGTGCTCCCTGCAATGTGATCTGCTCCATTATTTTCCAGAAACGTTTCGATTATAAAGATCAGCAATTTC
TTAACTTGATGGAAAAATTGAATGAAAACATCAGGATTGTAAGCACCCCCTGAATCCAGGTAAGGCCAAG
TTTTTTGCTTCCTGAGAAACCACTTACAGTCTTTTTTTCTGGGAAATCCAAAATTCTATATTGACCAAGC
CCTGAAGTACATTTTTGAATACTACAGTCTTGCCTAGACAGCCATGGGGTGAATATCTGGAAAAGATGGC
AAAGTTCTTTATTTTATGCACAGGAAATGAATATCCCAATATAGATCAGGCTTCTAAGCCCATTAGCTCC
```

**Figure S5.** Primary Sequence of IDT-Synthesized DNA Mutant-Type DNA template for CYP2C19*3

```
>NC_000010.11:g.94761746-94762123 Homo sapiens chromosome 10,
GRCh38.p14 Primary Assembly
TAAGTGGTTCTATTTAATGTGAAGCCTGTTTTATGAACAGGATGAATGTGGTATATATTCAGAATAACTA
ATGTTTGGAAGTTGTTTTGTTTTGCTAAAACAAAGTTTTAGCAAACGATTTTTTTTTTCAAATTTGTGTC
TTCTGTTCTCAAAGTATCTCTGATGTAAGAGATAATGCGCCACGATGGGCATCAGAAGACCTCAGCTCAA
ATCCCAGTTCTGCCAGCTATGAGCTGTGTGGCACCAACAGGTGTCCTGTTCTCCCAGGGTCTCCCTTTTC
CCATTTGAAATATAAAAAATAACAATTCCTGCCTTCACGTGTTTTTTTAGGGGGTTAAATGGTAAAGGTG
TTTATATCTGCTAAGGTAATTTACTTGA
```

**Figure S6.** Primary Sequence of IDT-Synthesized DNA Mutant-Type DNA template for CYP2C19*17

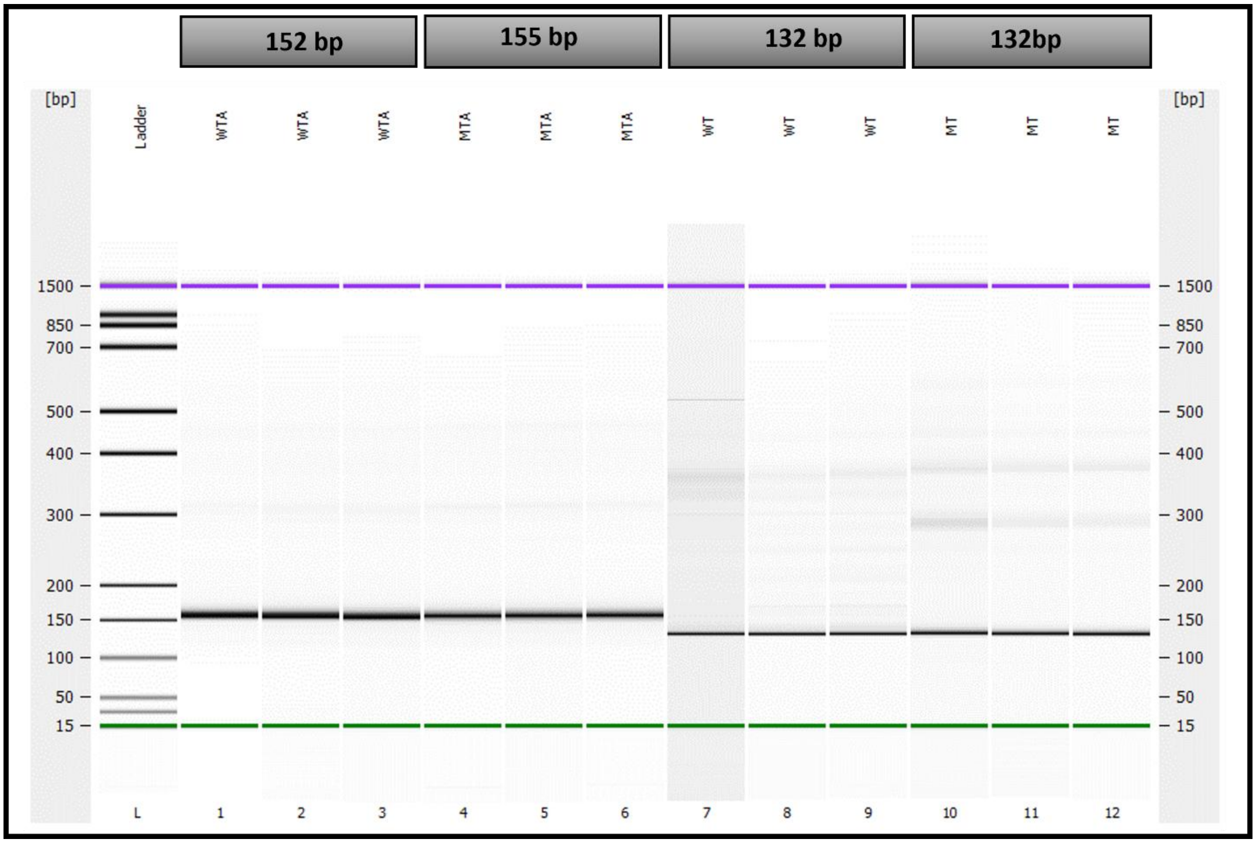


**Figure S7.** The selectivity of AS-PCR primers increases with the addition of adapter sequences. For the three replicates **with** adapter sequences that amplify the wild-type allele, *CYP2C19*1* (WTA), and the mutant-type allele, *CYP2C19*2* (MTA), single bands of 152bp and 155bp, respectively, were observed. This depicts high selectivity. However, for the three replicates **without** adapter sequences that amplify the wild-type allele, *CYP2C19*1* (WT), and the mutant-type allele, *CYP2C19*2* (MT), a prominent band of 132 bp was observed in both cases. However, stutter bands can be seen, which implies reduced primer selectivity.
